# SLC15A4 controls endolysosomal TLR7-9 responses by recruiting the innate immune adaptor TASL

**DOI:** 10.1101/2023.03.27.534109

**Authors:** Haobo Zhang, Léa Bernaleau, Maeva Delacrétaz, Ed Hasanovic, Hermann Eibel, Manuele Rebsamen

## Abstract

Nucleic acid sensing by endolysosomal Toll-like receptors (TLRs) plays a crucial role in innate immune responses to invading pathogens. In contrast, aberrant activation of these pathways is associated with several autoimmune diseases, such as systemic lupus erythematosus (SLE). The endolysosomal solute carrier family 15 member 4 (SLC15A4) is required for TLR7, TLR8 and TLR9-induced inflammatory responses and for disease development in different SLE models. SLC15A4 has been proposed to affect TLR7-9 activation through its transport activity, as well as by assembling in an IRF5-activating signalling complex with the innate immune adaptor TASL, but the relative contribution of these different functions remains unclear. Here we show that the essential role of SLC15A4 is to recruit TASL to the endolysosomal compartment, while its transport activity is dispensable. Targeting of TASL to the endolysosomal compartment is sufficient to rescue TLR7-9-induced IRF5 activation in SLC15A4-deficient cells. In line with this, lysosomal-localized TASL restored proinflammatory cytokines and type I interferon responses in absence of SLC15A4. Our study reveals that SLC15A4 acts as a signalling scaffold and that this transport-independent function is essential to control TLR7-9-mediated inflammatory responses. These findings further support targeting the SLC15A4-TASL complex as a potential therapeutic strategy for SLE and related diseases.

## INTRODUCTION

Detection of invading pathogens by the innate immune system is central to mount protective responses^1^. Microbial-derived nucleic acids are recognized by both cytosolic sensors as well as endolysosomal transmembrane Toll-like receptors (TLR) 3, 7, 8 and 9 ^2–5^. These innate immune pathways play a critical role to control viral and bacterial infections by inducing antimicrobial genes, triggering the production of interferons and proinflammatory cytokines and priming tailored adaptive immune responses. Conversely, aberrant activation of nucleic acid-sensing pathways is involved in a broad spectrum of pathologies, ranging from interferonopathies to autoimmune conditions such as systemic lupus erythematosus (SLE)^6–8^. A central pathogenic event in SLE and closely related autoimmune diseases is the engagement of endolysosomal TLRs, in particular TLR7, by endogenous, self-derived nucleic acids, resulting in the activation of immune cells, including primarily plasmacytoid dendritic cells (pDCs) and B cells^8–11^. These cells critically contribute to the development of the disease by producing type I interferons, proinflammatory cytokines and autoantibodies.

Over the past decade, the endolysosomal solute carrier family 15 member 4 (SLC15A4, also known as PHT1) has emerged as a critical component involved in TLR7-9-induced immune responses as well as in autoimmune diseases, a role strongly supported by both human genetics and animal studies. Indeed, evidences from genome-wide association studies (GWAS) implicated SLC15A4 in SLE^12–17^. The link between SLC15A4 and endosomal TLR7-9 responses was first revealed in an *in vivo* ENU mutagenesis screen assessing serum levels of type I IFNs upon injection of TLR7-9 agonists, which was impaired in *Slc15a4*-mutant *feeble* animals^18^. The requirement of this solute carrier for TLR7-9 function has been further established using conventional *Slc15a4*^-*/-*^ mice and by investigating different infections and autoimmune disease models, including chemically- and genetically-induced SLE^19–26^. These studies demonstrated that SLC15A4 deficiency impairs TLR7-9-induced responses in multiple cells types, comprising pDC and B cells, and confers significant protection to autoimmune diseases *in vivo*. Interestingly, beside TLR signalling, SLC15A4 has been implicated in other innate immune pathway, including NOD1-2 responses and inflammasome activation, and *Slc15a4* deficiency has been shown to be protective also in DSS-induced colitis models^24, 27–30^. Altogether, these studies support pharmacological inhibition of SLC15A4 as a potential therapeutic strategy for SLE and, possibly, other autoimmune and inflammatory conditions. Despite these findings, the mechanism(s) by which SLC15A4 affects TLR7-9 responses remains less clear, and multiple explanations have been proposed. The SLC15A family comprises five members (1-5) and the best characterized, SLC15A1 (PepT1) and SLC15A2 (PepT2), act as plasma membrane proton-coupled oligopeptide transporters^31^. Similarly, SLC15A4 has been described as an endolysosomal proton-coupled transporter mediating histidine/oligopeptide translocation from the lumen to the cytosol^21, 30, 32–34^. Based on this function, SLC15A4 deficiency has been proposed to impair TLR7-9 function by altering endolysosomal homeostasis, pH and/or histidine concentration influencing thereby TLR maturation, TLR-ligand engagement, mTORC1 activity or cellular metabolic processes^18, 21, 24, 33, 35, 36^. Furthermore, it was recently advanced that SLC15A4 deficiency compromises the trafficking of TLRs and their ligands to endolysosomes leading to defects in receptor engagement and in the generation of an endolysosomal organelle required for efficient signalling^23^.

Investigating this critical mechanistic aspect, we recently showed that SLC15A4 forms a signalling complex with a previously uncharacterized protein encoded on the chromosome X by the SLE-associated gene *CXorf21*, which we named TASL^37, 38^. Loss of TASL or mutations impairing complex formation phenocopied SLC15A4 deficiency, resulting in impaired type I IFN and proinflammatory cytokine production upon TLR7-9 stimulation^37^. Importantly, both SLC15A4 or TASL knockout specifically impaired TLR7-9-induced IRF5 activation without affecting NF-kB and MAPK pathway, strongly suggesting that TLR-ligand engagement still occurs in SLC15A4- and TASL-deficient cells and that this complex specifically affects TLR signalling downstream of this initiating event. In line with this, we identified in the C-terminal region of TASL a pLxIS motif, which in the IRF3-adaptor proteins MAVS, STING and TRIF is required for IRF3 recruitment and activation^39^. Analogously, TASL pLxIS motif was essential for IRF5 binding, phosphorylation and downstream transcriptional responses^37^. Altogether, our study revealed that SLC15A4 controls IRF5 activation by mediating the recruitment to the endolysosomal compartment of TASL, which, through its pLxIS motif, acts as a novel IRF5-activating immune adaptor^37^.

Collectively, these studies raise the question of the relative contribution for endolysosomal TLR7-9 responses of the two proposed functions of SLC15A4, i.e. transporter and TASL-recruiting signalling complex. Indeed, our data showed that the transport-inactivating mutations E465K/A, previously used to demonstrate the importance of the SLC15A4 transporter function^21^, also resulted in a complete impairment of TASL binding^37^. A detailed mechanistic understanding of the role of SLC15A4 is key to further evaluate its potential as drug target for SLE and inform efforts aiming at pharmacologically interfering with its function. Here we show that the critical role of SLC15A4 in endolysosomal TLR7-9 responses is to mediate lysosomal recruitment of TASL, and that SLC15A4-mediated transport activity is dispensable for IRF5 activation and proinflammatory responses.

## RESULTS

### Fusion of TASL to transport-deficient SLC15A4(E465A) rescues TLR7/8 responses

In order to assess the relative contribution of SLC15A4 transport activity and endolysosomal TASL recruitment for TLR-induced responses, we devised a strategy to uncouple these two functions by fusing TASL coding sequence to the cytoplasmic C-terminus of SLC15A4, either wildtype or bearing the E465A substitution (Figure 1A). Mutations of the key transmembrane residue E465 in SLC15A4 have been shown to impair both its transport function as well as TASL binding^21, 37^. Accordingly, SLC15A4 E465 mutants failed to rescue TLR7-9-induced signalling when expressed in SLC15A4-deficient cells^37^. We first verified that fusion of TASL to SLC15A4 C-terminus did not alter its trafficking to the endolysosomal compartment. SLC15A4-TASL and SLC15A4(E465A)-TASL fusion proteins showed the expected glycosylation when stably expressed in human monocytic THP1 cells (Figure S1A). In line with this, both fusion proteins were detected on the LAMP2-positive lysosomes in these TLR7/8-signalling competent cells (Figure S1B).

**Figure 1.**
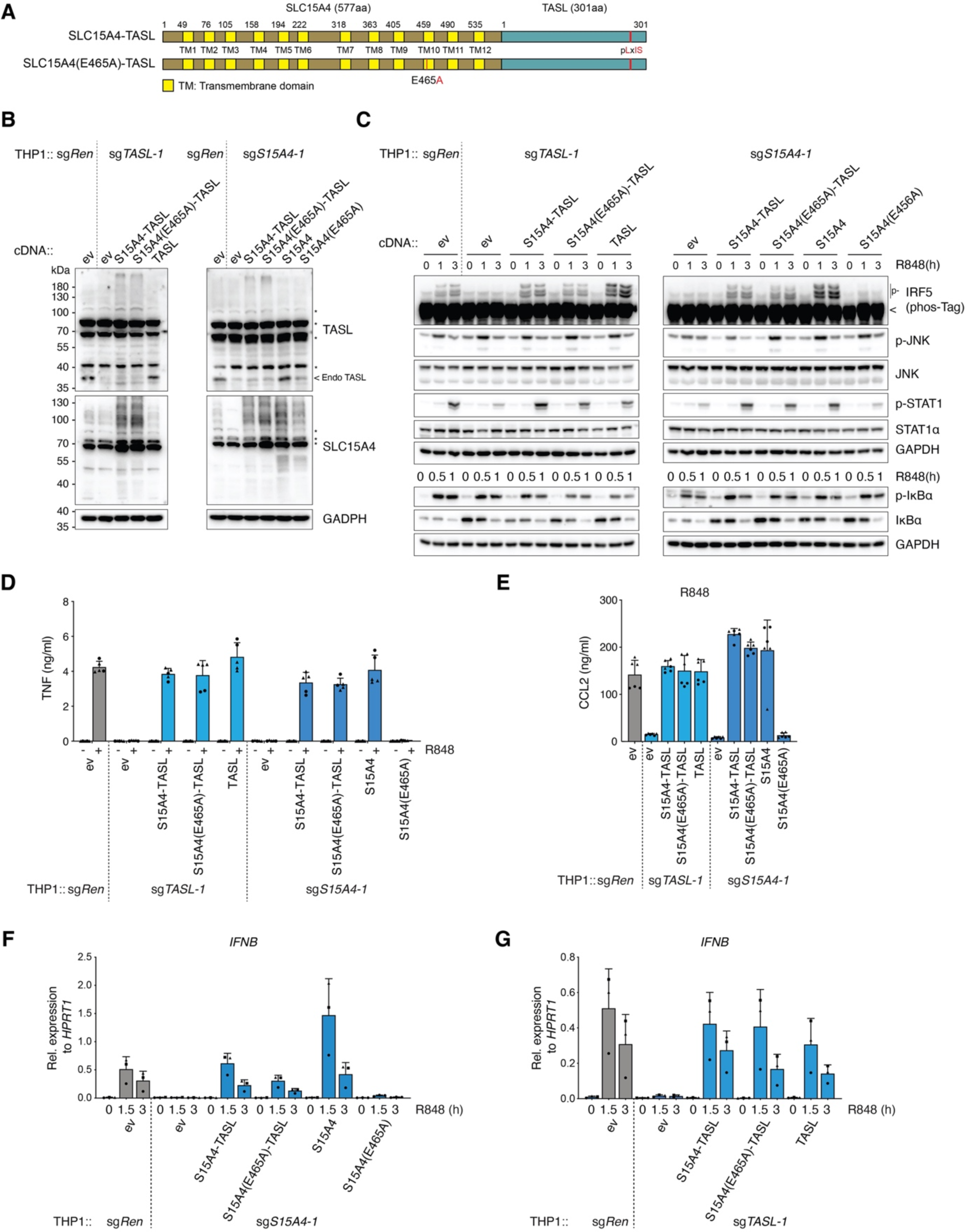
SLC15A4 transporter activity is dispensable for TLR7/8-induced IRF5 signalling. (A) Schematic of SLC15A4-TASL and SLC15A4(E465A)-TASL fusion proteins. (B) Immunoblots of THP1 cells carrying sgRNAs targeting *SLC15A4* (sg*SLC15A4*-1), *TASL* (sg*TASL*-1) or a control sgRNA targeting *Renilla* (sg*Ren*) and stably expressing the indicated constructs. Cell lysates were subjected to immunoblots with the indicated antibodies. ev: empty vector. Asterisks represent unspecific bands. Arrow indicates endogenous TASL. (C) Immunoblots of lysates from the indicated THP1 cell lines stimulated with R848 (5 μg ml−1 for 0-3 h). p-: phosphorylated. Phos-Tag: phos-Tag-containing gel. (D and E) Cytokine production of indicated THP1 cells unstimulated or stimulated with R848 (5 μg ml−1 for 24h). (F and G) *IFNB* mRNA levels of indicated THP1 cells stimulated with R848 (5 μg ml−1 for 0-3 h) measured by quantitative real-time PCR (qPCR). In B and C, data are representative of two independent experiments. In D and E, graphs show mean ± s.d. (n = 2 biologically independent experiments performed in stimulation replicates are shown). In F and G, mean ± s.d. (n = 3 biological replicates).

Next, we stably expressed these constructs in SLC15A4-or TASL-deficient THP1 cells to assess their activity (Figure 1B). Of note, TASL protein stability depends on SLC15A4 binding^37^, resulting in the lower endogenous levels observed in SLC15A4 knockout lines. As expected, expression of wildtype SLC15A4 but not SLC15A4(E465A) in SLC15A4 knockout cells restored IRF5 activation upon stimulation with TLR7/8 agonist R848 (resiquimod), as assessed by IRF5 phosphorylation on phos-Tag-containing gels (Figure 1C). Importantly, SLC15A4-TASL and, most notably, SLC15A4(E465A)-TASL constructs fully rescued IRF5 activation in SLC15A4-deficient cells, indicating that fusion of TASL to the inactive SLC15A4(E465A) restored its functionality (Figure 1C). These data were further confirmed monitoring IRF5 phosphorylation and dimerization upon stimulation with the TLR8-specific agonist TL8-506 (Figure S1C-D). Similar results were obtained when we assessed these constructs in TASL-knockout THP1 cells, with both SLC15A4-TASL and SLC15A4(E465A)-TASL supporting TLR7/8-induced IRF5 activation (Figure 1C). Activation of STAT1, likely resulting from IFN paracrine signalling, was equally restored by SLC15A4-TASL fusion proteins (Figure 1C). Finally, NF-kB and MAPK-pathway activation, monitored by IkBα and JNK phosphorylation respectively, proceeded independently of SLC15A4 and TASL, confirming that this complex acts downstream of TLR-ligand engagement to specifically control the IRF5 signalling branch (Figure 1C).

Next, we investigated whether SLC15A4-TASL and, in particular, SLC15A4(E465A)-TASL fusions retained the full spectrum of SLC15A4 and TASL activities by assessing the effect on downstream inflammatory cytokines and chemokines production. Expression of SLC15A4-TASL and SLC15A4(E465A)-TASL rescued both TNF and CCL2 production in SLC15A4-deficient THP1, while SLC15A4(E465A) had no effect as expected (Figure 1D-E). In line with this (and correlating with STAT1 activation), *IFNB* induction upon R848 stimulation was restored in SLC15A4 knockouts expressing either SLC15A4-TASL or SLC15A4(E465A)- TASL, but not SLC15A4(E465A) (Figure 1F). Interestingly, both SLC15A4-TASL fusions normalized cytokine/chemokine production and *IFNB* induction also in TASL knockout cells, suggesting that TASL does not need to be released from the endolysosomal compartment to fulfil its function (Figure 1D-E and 1G).

Altogether, these results strongly suggest that SLC15A4 transport activity is not essential for TLR7/8-induced IRF5 activation and downstream signalling. Rather, they support the notion that the crucial role of this solute carrier in endolysosomal TLR pathway is to act as a signalling complex mediating the recruitment of TASL to the endolysosomal compartment.

### Endolysosomal targeted TASL sustains TLR7/8-induced IRF5 activation independently of SLC15A4

These findings raised the question whether endolysosomal TASL localization is in itself sufficient to mediate TLR7/8-induced IRF5 activation independently of any other possible SLC15A4 function(s). To assess this, we targeted TASL to the lysosomal compartment independently of SLC15A4 by generating fusion constructs with LAMP1 or LAMTOR1, the lysosomal-anchoring component of the Ragulator complex (Figure 2A)^40^. We selected these two lysosomal proteins because they allow to anchor TASL to this compartment using a different mechanism than the multitransmembrane SLC15A4. In the case of LAMP1, TASL sequence was inserted after the short, 12 amino acid-long cytoplasmic sequence which follows its single transmembrane domain. In contrast, LAMTOR1 does not contain any transmembrane domains and its lysosomal localization is mediated by N-terminal lipidation (myristoylation of Gly2 and palmitoylation of Cys3 and Cys4)^40^. LAMTOR1 N terminus has been previously shown to be sufficient for lysosomal localization, we therefore generated two different TASL fusion constructs, containing the first 1-39 or 1-81 amino acids of LAMTOR1^40^. Of note, both constructs do not contain the LAMTOR1 region required for binding to the other subunits of the Ragulator complex (LAMTOR2-5), minimizing therefore any risk of interfering with its functions^40–42^. Upon stable expression in THP1 cells, LAMP1-TASL and LAMTOR1-TASL fusions were targeted to the lysosomal compartment (Figure 2B-C, Figure S2A). We have previously shown that the first, N-terminal amino acids of TASL are required for SLC15A4 binding and that TASL N-terminal tagging impaired complex formation^37^. Therefore, we first investigated whether fusion of the lysosomal targeting proteins to TASL N-terminus would affect SLC15A4 binding. Indeed, TASL fusions failed to co-immunoprecipitate SLC15A4 when co-expressed in HEK293T cells (Figure S2B). In line with this, immunoprecipitation of TASL fusion proteins stably expressed in THP1 cells did not recover endogenous SLC15A4 (Figure S2C). Altogether, these results indicate that LAMTOR1-and LAMP1-TASL localized to the lysosomal compartment independently of SLC15A4.

**Figure 2.**
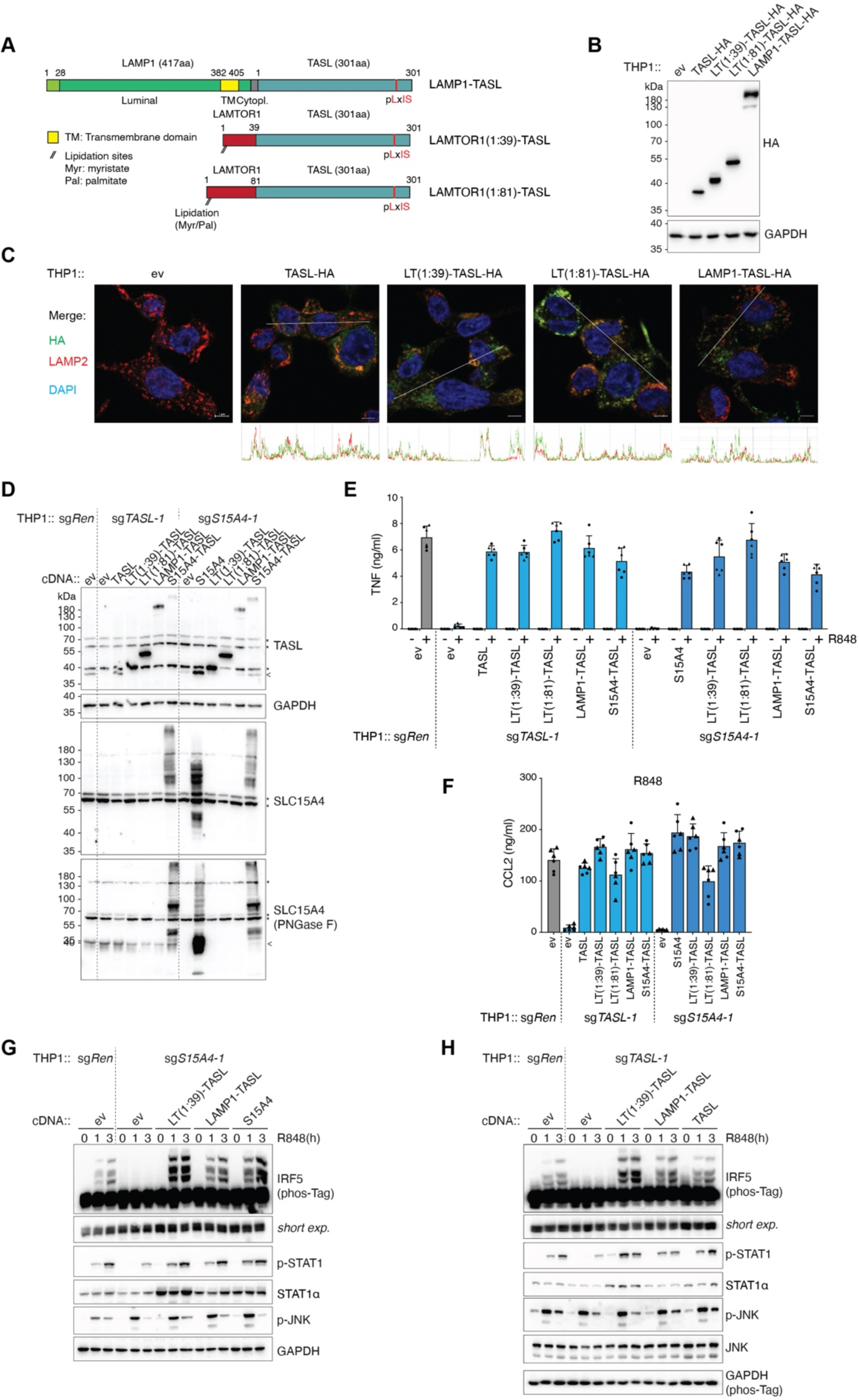
Endolysosomal targeted TASL rescues TLR7/8-induced IRF5 signalling independently of SLC15A4. (A) Schematic of LAMP1-TASL, LAMTOR1(1:39)-TASL and LAMTOR1(1:81)-TASL fusion proteins. TM: Transmembrane domain; Myr: myristate; Pal: palmitate. (B) Immunoblots of THP1 cells stably expressing the indicated C-terminal HA-tagged TASL fusion proteins. (C) Confocal microscopy of indicated THP1 cells. Green: anti-HA; red: anti-LAMP2; blue: DAPI. Scale bar: 5 μm (D) Immunoblots of lysates from knockout THP1 cells stably reconstituted with indicated constructs. Where indicated, lysates were treated with PNGase F. Asterisk represents an unspecific band. Arrows indicates endogenous proteins. (E) TNF production of indicated THP1 cells unstimulated or stimulated with R848 (5 μg ml−1 for 24h). (F) CCL2 production of indicated THP1 cells treated with R848 (5 μg ml−1 for 24h). (G and H) Immunoblots of lysates from the indicated THP1 cells stimulated with R848 (5 μg ml−1 for 0-3 h). p-, phosphorylated. In B, D, G and H, data are representative of two independent experiments. In E and F, mean ± s.d. (n = 2 biologically independent experiments performed in stimulation replicates are shown).

Next, we stably expressed these fusion constructs in SLC15A4- and TASL-deficient THP1 cells, along with SLC15A4-TASL and the respective wildtype controls (Figure 2D). Remarkably, the two LAMTOR1-TASL constructs as well as LAMP1-TASL fully normalized R848-induced TNF production in both knockout cell lines (Figure 2E). CCL2 production was equally restored (Figure 2F). Supporting these data, expression of lysosomal-localized TASL in SLC15A4-deficient cells efficiently rescued IRF5 activation, monitored by its phosphorylation and dimerization, as well as STAT1 phosphorylation (Figure 2G, Figure S2D). Consistently, LAMTOR1- and LAMP1-TASL efficiently complemented TLR7/8 signalling also in TASL knockout cells (Figure 2H). Of note, compared to control sgRen cells, lysosomal TASL fusions, and especially LAMTOR1(1:39)-TASL, showed a mild increase in R848-induced IRF5 activation. This may be due to higher expression levels compared to endogenous TASL levels or, possibly, by its stable lysosomal association. To complement these data obtained upon stable expression of TASL fusion constructs, we generated SLC15A4-deficient cells inducibly expressing LAMTOR1(1:39)-TASL upon doxycycline treatment, allowing to transiently express this construct to levels comparable with endogenous TASL protein (Figure S2E). LAMTOR1(1:39)-TASL restored IRF5 activation also in these conditions, indicating that lysosomal targeting of TASL in absence of SLC15A4 can achieve comparable activity as the endogenous protein in wildtype cells (Figure S2E). Altogether, these results reveal that the key function of SLC15A4 essential for TLR7/8-induced responses is its ability to recruit TASL to lysosomes, as lysosomal localization of TASL is sufficient to fully restore IRF5 activation and downstream cytokine/chemokine production in SLC15A4-deficient cells. Therefore, SLC15A4-mediated transport and/or other direct or indirect metabolic effects appear to be dispensable for TLR7/8-induced IRF5-dependent responses.

### Endolysosomal TASL restores TLR7-9 responses in SLC15A4-deficient pDC and B cells

To further confirm these findings, assess possible cell-type specific effects, and explore TLR9-induced responses, we next investigated the human plasmacytoid dendritic cell line CAL-1^43^. Plasmacytoid DCs are major producers of type I IFN production upon endolysosomal TLR stimulation by microbial or endogenous nucleic acids and play a central role in SLE pathogenesis^8, 44^. Moreover, CAL-1 cells express TLR9 and we previously showed that stimulation with its ligand CpG triggers SLC15A4- and TASL-dependent IRF5 activation, allowing therefore to extend our investigation beyond TLR7/8^37^. Consistently with the data obtained in THP1 monocytes, stable expression of LAMTOR1-, LAMP1- and SLC15A4-TASL fusion constructs restored R848-induced IRF5 activation in both SLC15A4- and TASL-deficient CAL-1 cells (Figure 3A-C, Figure S3A-B). Importantly, the capacity of lysosomal-targeted TASL to rescue IRF5 activation in absence of SLC15A4 was not specific to TLR7/8, but also observed upon TLR9 stimulation. Indeed, the impaired IRF5 activation observed in SLC15A4-or TASL-knockout CAL-1 upon stimulation with TLR9 agonist CpG was largely restored by expression of lysosomal TASL fusion proteins (Figure 3D-E, Figure S3C-D). Similar effects were observed when monitoring STAT1 activation, while, as expected, MAPK pathway proceeded independently of SLC15A4-TASL complex, confirming unaltered TLR-ligand engagement. Mirroring IRF5 activation, TNF and IL6 production upon R848 and CpG was efficiently rescued by lysosomal TASL in both CAL-1 knockout lines (Figure 3F-G).

**Figure 3.**
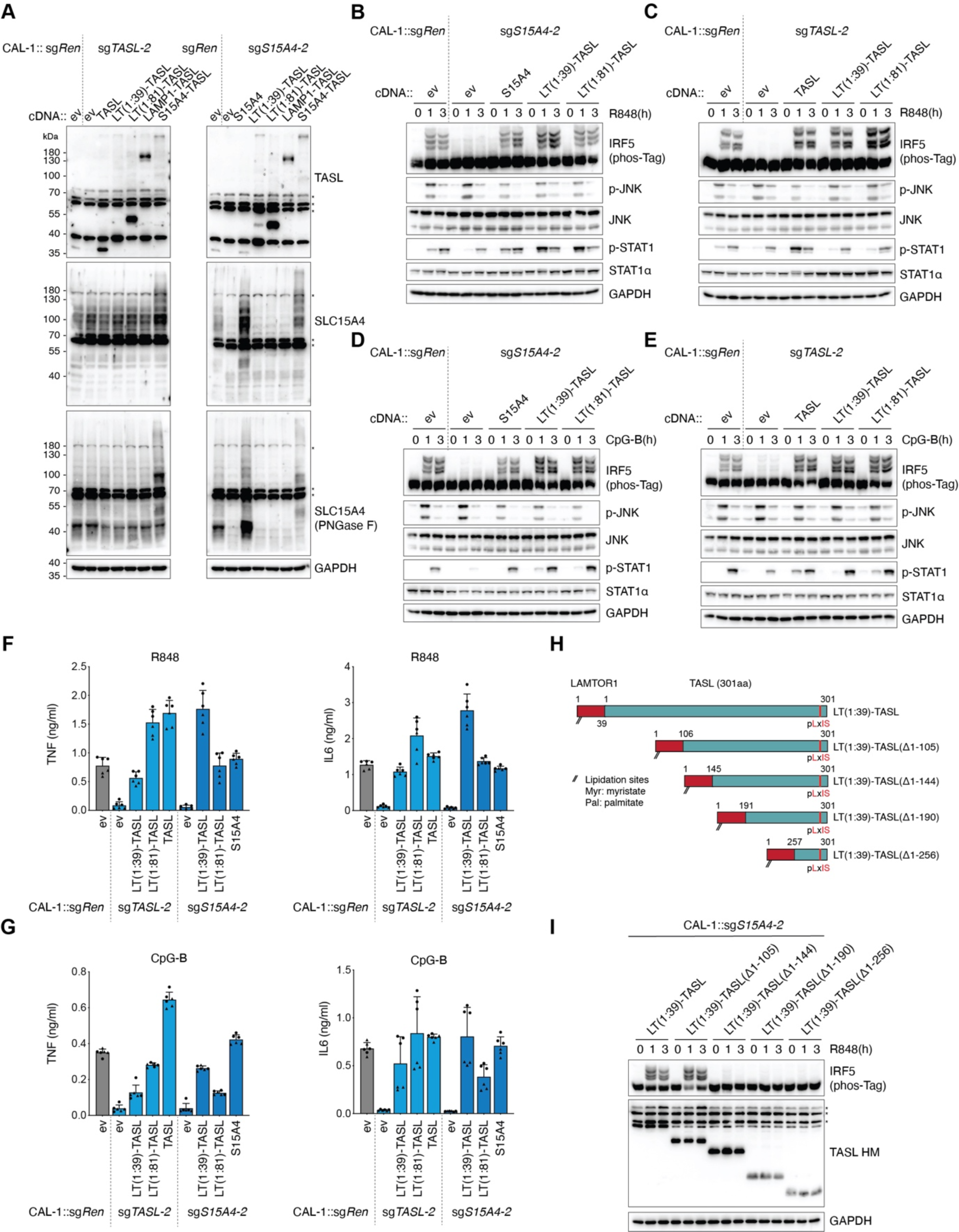
Endolysosomal TASL is sufficient to restore TLR7/8 and TLR9 responses in SLC15A4-deficient CAL-1 pDCs. (A) Immunoblots of knockout CAL-1 cells stably reconstituted with indicated constructs. Where indicated, lysates were treated with PNGase F. Asterisks denote unspecific bands. (B-E) Immunoblots of lysates from the indicated CAL-1 cells stimulated with R848 (5 μg ml−1 for 0-3 h) (B and C) or CpG-B (5 μM for 0-3 h) (D and E). (F and G) TNF (left) and IL6 (right) production of indicated CAL-1 cells stimulated for 24h with R848 (5 μg ml−1) (F) or CpG-B (5 μM) (G). (H) Schematic of LAMTOR1(1:39)-TASL deletion constructs. (I) Immunoblots of indicated CAL-1 cells stimulated with R848 (5 μg ml−1 for 0-3 h). Asterisks denote unspecific bands. In A-E and I, data are representative of two independent experiments. In F and G, graphs show mean ± s.d. (n = 2 biologically independent experiments performed in stimulation replicates are shown).

Next, we took advantage of the fact that lysosomal TASL fusions allow to uncouple localization and signalling activity to identify domains in TASL specifically required for IRF5 activation. For this we expressed a series of TASL deletion constructs bearing the lysosomal LAMTOR1(1:39)-targeting motif in SLC15A4-deficient cells (Figure 3H-I, Figure S3E). Interestingly, while lysosomal TASL deleted of amino acids 1-105 retained full activity, further removal of the following evolutionary conserved region (aa 106-144) resulted in a complete loss of IRF5 phosphorylation, suggesting a third functional motif in TASL in addition to the N-terminal lysosomal targeting region and C-terminal IRF5-recruiting pLxIS motif. Of note, stable expression of wildtype TASL in SLC15A4 knockout cells was unable to restore IRF5 activation, further confirming that lysosomal anchoring, and not increased TASL expression levels, is crucial for restoring signalling upon loss of this solute carrier (Figure S3F).

Besides pDCs, B cells are key contributors to autoimmune disease pathogenesis^9, 45^. In this context, accumulating evidences support a central role of cell-intrinsic endolysosomal TLR signalling in promoting the main processes by which B cells contribute to autoimmune diseases, including production of autoantibodies, antigen presentation to T cells and production of cytokines^9, 11^. *Slc15a4*-deficiency in B cells affects their function and confers protection in murine SLE models^21, 25, 26^, but the role of this solute carrier in human B cells has not been yet investigated. Moreover, the function of TASL has not been explored in neither murine nor human B cells. We therefore assessed the involvement of the SLC15A4-TASL complex for endolysosomal TLR responses and IRF5 activation in EBV-immortalized human B cell lines. After verifying that R848 stimulation triggered IRF5 activation in two independent lines, we generated SLC15A4 and TASL knockout (Figure S4A, S4C). In line with previous observations in THP1 and CAL-1 cells (Figure 1B, 3A)^37^, deletion of SLC15A4 resulted in a concomitant reduction in TASL protein levels, suggesting functional complex formation also in these cells (Figure S4A, S4C). Knockout of either SLC15A4 or TASL strongly impaired IRF5 phosphorylation and dimerization in both lines, with the reduction in IRF5 activation correlating with the knockout efficiency of the different sgRNAs observed in these cell populations (Figure 4A-B, Figure S4B, S4D-E). These data demonstrate that the SLC15A4-TASL complex is essential for endolysosomal TLR-induced IRF5 activation also in human B cells, extending therefore its role beyond innate immune monocytes and pDCs, and demonstrating its central relevance in the main TLR7-9 responding cell types. Lastly, we assessed whether lysosomal targeting of TASL in absence of SLC15A4 was sufficient for IRF5 activation. Consistently with results obtained in monocytic and pDC cells, lysosomal-localized TASL rescued IRF5 activation in SLC15A4- and TASL-deficient cells to a level comparable to the cells reconstituted with respective wild-type proteins (Figure 4C-F). Considering the central role of both B cells and IRF5 in autoimmune conditions and SLE in particular, these results further emphasize the therapeutic potential of targeting the SLC15A4-TASL complex in these diseases.

**Figure 4.**
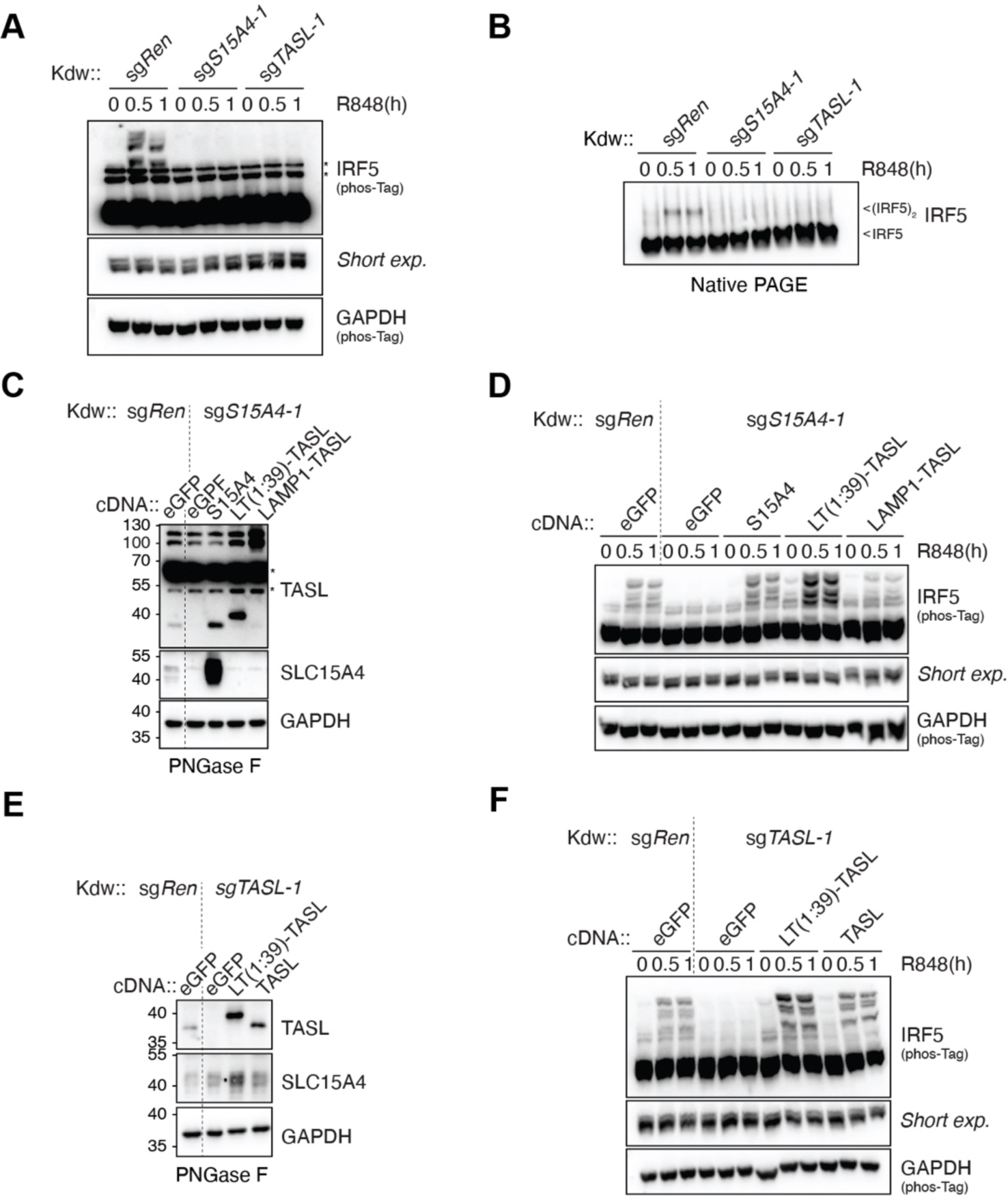
SLC15A4-TASL complex, independently of SLC15A4 transporter activity, is essential for endolysosomal TLR-induced IRF5 activation in human B cells. (A) Immunoblots of indicated knockout Kdw cells stimulated with R848 (5 μg ml−1, for 0-1h). Asterisks denote unspecific bands. (B) Native PAGE immunoblot analysis of IRF5 in knockout Kdw cells stimulated with R848 (5 μg ml−1, for 0-1h). Arrows indicate IRF5 monomer or dimer. (C-F) Immunoblots of SLC15A4 (C and D) or TASL knockout Kdw cells stably reconstituted with indicated constructs in unstimulated (C and E) or stimulated (D and F) conditions (R848, 5 μg ml−1, for 0-1h). In (C and E) lysates were treated with PNGase F. eGFP: enhanced GFP. Asterisks denote unspecific bands. In A-F, data are representative of two independent experiments.

## DISCUSSION

In this study, we show that the crucial role of SLC15A4 in controlling TLR7-9-induced IRF5 activation and cytokine production is to recruit TASL to the endolysosomal compartment, while its substrate transport activity is not required *per se*.

How SLC15A4 controls endolysosomal TLR7-9 responses remained unclear and different mechanistic explanations, not all necessarily mutually exclusive, have been proposed. Early studies support a model in which loss of SLC15A4 proton-coupled histidine/oligopeptide transport activity results in the accumulation of its substrates in the endolysosomal lumen, altering thereby pH and/or histidine levels^18, 21, 24, 36^. This in turn would impact on TLR maturation and/or ligand-receptor engagement. Altered intralysosomal environment and consequent impaired mTORC1 activation have been also proposed to influence TLR responses by affecting a feedforward loop mediating IRF7 upregulation^21^. Finally, other studies linked SLC15A4-deficiency to perturbed trafficking and colocalization of TLRs with their ligand, resulting in impaired receptor-ligand engagement^23^, or to metabolic perturbations affecting the TCA cycle, autophagy and/or mitochondrial integrity^33, 35^. The finding that lysosomal localization of TASL is sufficient to rescue TLR7-9 responses in SLC15A4-deficient cells demonstrates that translocation or altered concentrations of potential substrates of SLC15A4 is not critically involved in TLR pathway regulation, at least in terms of IRF5 activation and production of proinflammatory cytokines and Type I IFNs. Importantly, this does not exclude the possibility that SLC15A4 transport activity may indirectly affect the TLR7-9 pathway by controlling TASL recruitment. Indeed, it is conceivable that transport-dependent conformational changes in SLC15A4 could impact its ability to bind TASL, and therefore indirectly regulate IRF5 activation and downstream signalling. Whether the ability of SLC15A4 to recruit TASL is dependent on its conformation and/or its localization along the endolysosomal system is an intriguing question to be addressed in future studies.

Concerning the function of SLC15A4 as transporter, it should be noted that this solute carrier is expressed broadly, including in cell types that do not express TLR7-9, TASL nor IRF5^37^. This strongly suggests that SLC15A4 is involved in other cellular processes beside its specific TASL-recruiting function in endolysosomal TLR signalling. This is consistent with its reported function in regulating cellular metabolism, mTORC1 activation, NOD-ligand transport and mast cell responses^24, 27–30, 35, 46^.

The involvement of endosomal TLR responses in SLE and related diseases is strongly supported by both animal studies and human genetics. Notably, the three components of the signalling axis we described, SLC15A4, CXorf21/TASL and IRF5, have all been identified in GWAS on human SLE, with IRF5 being one of the best characterized and strongly associated factors^12, 13, 47, 48^. In line with this, IRF5-deficient mice show strong protection in a broad range of SLE disease models^49–54^. These evidences and the fact that solute carriers are an eminently druggable class of proteins, have put forward SLC15A4 as an attractive drug target for SLE and related diseases^55–57^. Further supporting this notion, here we show that the SLC15A4-TASL complex is essential for IRF5 activation not only in monocytes and pDCs, but also in human B cells, demonstrating therefore its general requirement in all the endolysosomal TLR-responding cells shown to be critically involved in SLE pathogenesis. The data presented strongly suggest that future efforts to pharmacologically target SLC15A4 should aim at interfering with SLC15A4-TASL complex function, either by direct inhibition of its assembly or, possibly, indirectly by interfering with the trafficking of the complex to the endolysosomal compartment. In contrast, our data imply that inhibition of SLC15A4 transport activity may not be in itself sufficient to inhibit endolysosomal TLR responses if this does not concomitantly result in interfering with complex formation or localization.

Finally, from the perspective of understanding the solute carrier family and its biology, our findings further highlight the fact that SLCs can have, behind their canonical role as transporters, additional, still underappreciated functions by acting as transceptor (transporter-receptor) and signalling complexes^58, 59^.

## Acknowledgments

We would like to thank Ales Drobek and the members of the Rebsamen laboratory for discussions and suggestions; Giulio Superti-Furga (CeMM-Center for Molecular Medicine) for kindly providing critical reagents and Takahiro Maeda (Nagasaki University) for sharing CAL-1 cells, Laurence Romy and Margot Thome-Miazza (University of Lausanne) for experimental suggestions, Leonhard Heinz (Medical University of Vienna) for critically reading the manuscript, and the Cellular Imaging Facility of the UNIL for technical support. This work was supported by the Swiss National Science Foundation (Project grant 310030_200709).

## Authors contributions

Conceptualization: HZ, MR; Data curation: HZ, LB, MR; Formal Analysis: HZ, LB, MD, MR; Funding acquisition: MR; Investigation: HZ, LB, MD, EH, MR; Methodology: HZ, LB, HE, MR; Project administration: MR; Resources: EH, MR; Supervision: MR; Validation: HZ, LB, MD, MR; Visualization: HZ, LB, MD, EH, MR; Writing – original draft: MR; Writing – review & editing: HZ, LB, MD, EH, HE, MR.

## Declaration of interests

M.R has filed patent applications related to the SLC15A4-TASL complex. The other authors declare no competing interests.

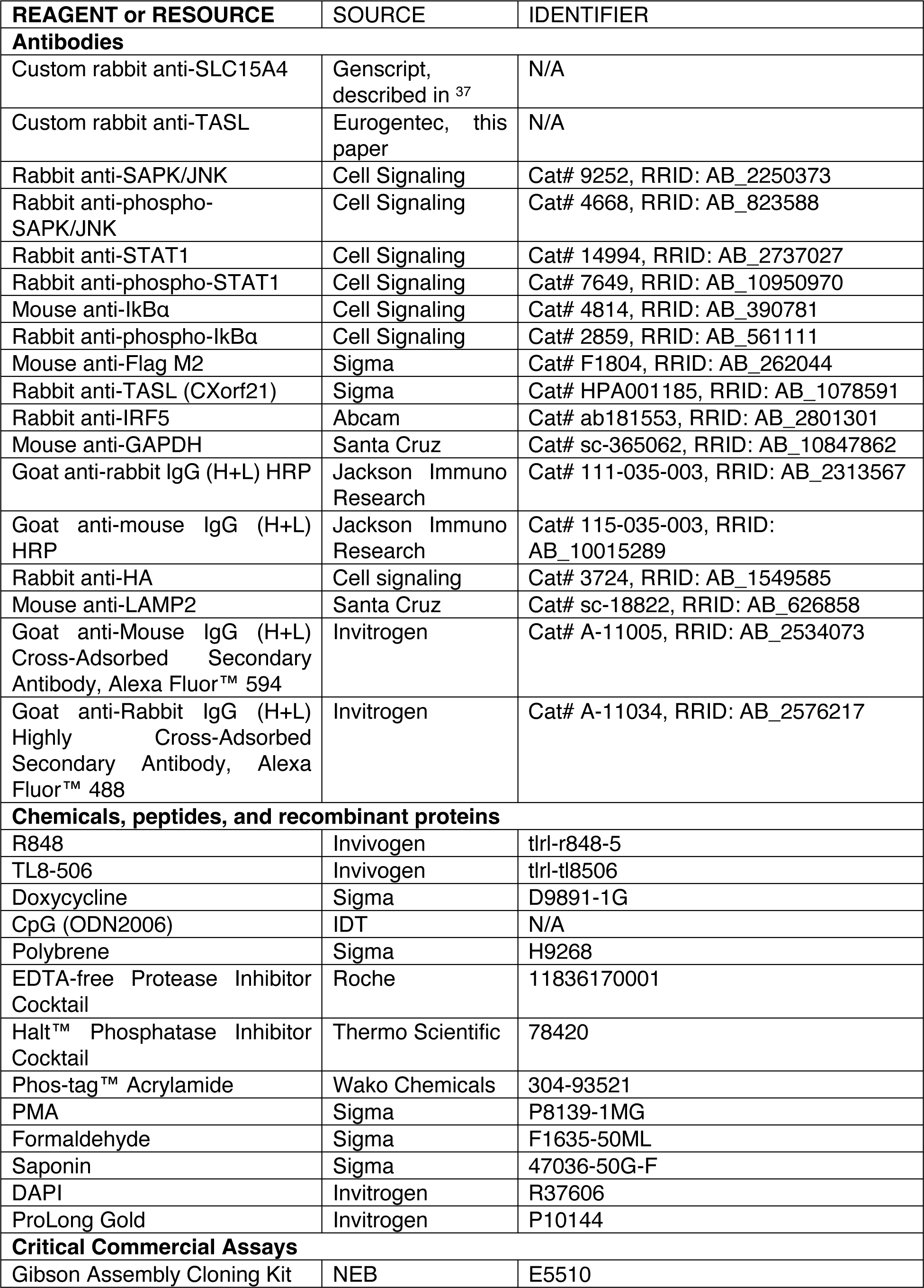
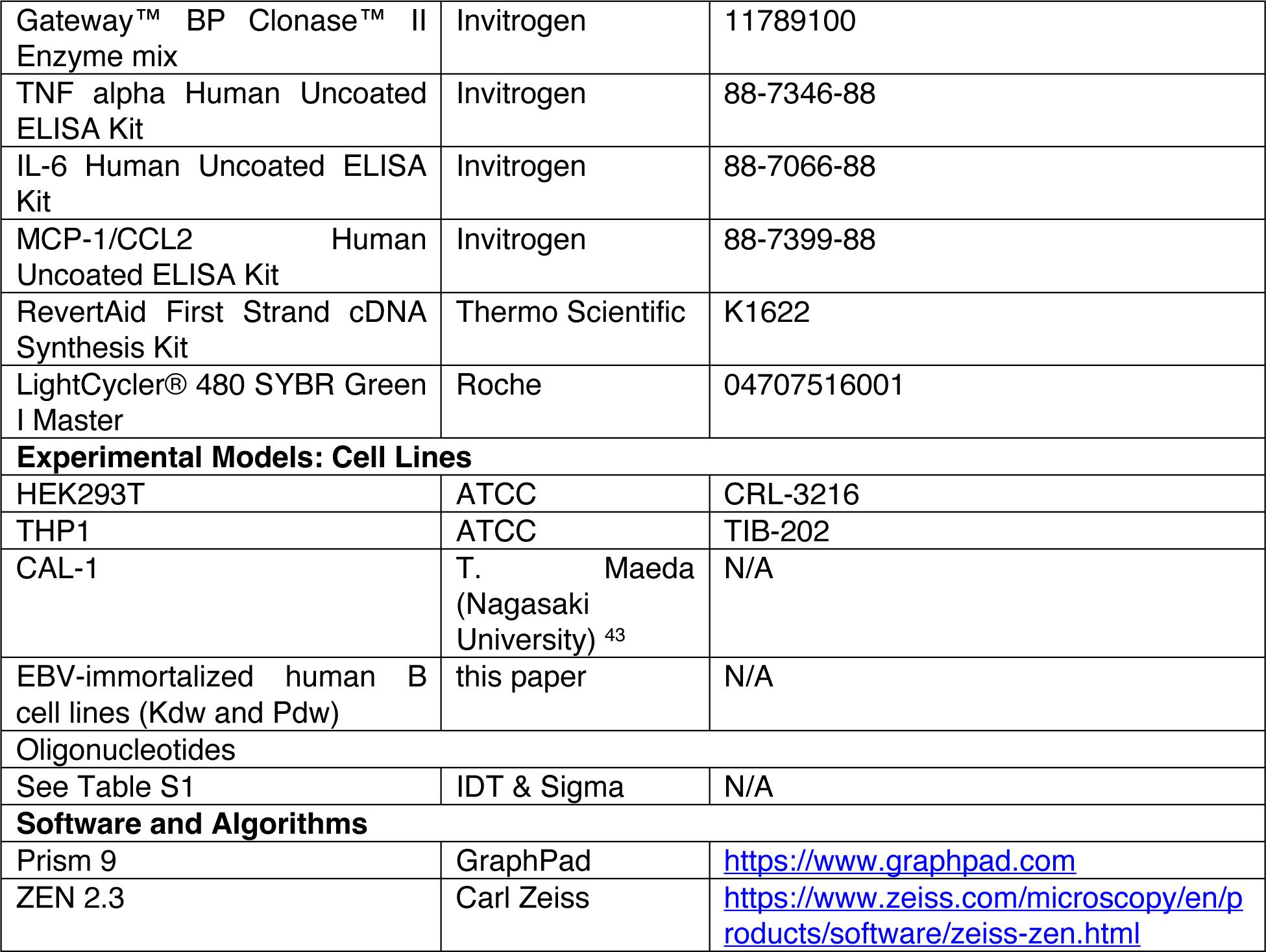
Key resources table

## Resource availability

### Lead contact

Further information and requests for resources and reagents should be directed to and will be fulfilled by the Lead Contact, Prof. Manuele Rebsamen.

### Materials availability

All unique reagents generated in this study are available from the lead contact upon completion of a Material Transfer Agreement.

### Data and code availability

This study did not generate any unique datasets or code.

## Experimental model and subject details

### Cells

HEK293T cells and THP1 cells were purchased from ATCC. CAL-1 cells were kindly provided by T. Maeda (Nagasaki University)^43^. EBV lines (Kdw and Pdw) were obtained by immortalizing B cells of healthy donors as described in ^60^. HEK293T cells were cultured in DMEM (Gibco), THP1, CAL-1 and B cells in RPMI (Gibco), supplemented with 10% (v/v) fetal bovine serum (FBS, Gibco) and antibiotics (100 U/ml penicillin, 100 μg/ml streptomycin, Bioconcept) at 37 °C in 5% CO2 incubator.

### Plasmids

Codon-optimized cDNAs for human SLC15A4 and TASL have been described previously ^37^. A template for cloning human LAMP1 (no. 134868) and DOX-inducible lentiviral gene expression vector pINDUCER21 (ORF-EG) (no. 46948) were from Addgene. Gateway pDONR201 plasmid LAMTOR1 has been described before ^59^. Gibson assembly cloning (NEB) was used to clone endolysosomal TASL fusion constructs (SLC15A4-TASL, SLC15A4(E465A)-TASL, LAMTOR1(1:39)-TASL, LAMTOR1(1:81)-TASL and LAMP1-linker(13 aa)-TASL (referred to as LAMP1-TASL)). Point mutation SLC15A4(E465A) was introduced by site-directed mutagenesis. LAMTOR1(1:39)-TASL deletion constructs (LAMTOR1(1:39)-TASL Δ1-105, Δ1-144, Δ1-190, Δ1-256) were generated by PCR mutagenesis. C-tag or HA-tag was added to the C-terminus of different TASL fusions by PCR. All cDNAs were subcloned into pDONR201/221 (Invitrogen), verified by sequencing and shuttled by Gateway cloning (Invitrogen) to destination vectors: pINDUCER21 or pRRL-based lentiviral expression plasmids with a selectable resistance cassette allowing untagged, N-or C-terminal Strep-HA-tagged (SH) or V5-tagged expression ^37^. CRISPR–Cas9-based knockout cell line generation was performed using pLentiCRISPRv2 (Addgene plasmid no. 52961) and sgRNA sequences targeting *SLC15A4* (sg*SLC15A4*-1 and sg*SLC15A4*-2), *TASL* (sg*TASL*-1 and sg*TASL*-2) or a non-targeting, control sgRNA sequence designed against *Renilla* (sg*Ren*) previously described (Table S1) ^37^.

## Method details

### Generation of stable knockout and overexpressing cell lines by lentiviral transduction

Lentiviral transduction was performed as previously described ^37^. Briefly, HEK293T cells were transfected with sgRNA-or cDNA-encoding lentiviral vectors and packaging plasmids psPAX2 and pMD2.G (plasmid no. 12260 and plasmid no. 12259 from Addgene) using PEI (Sigma). The medium was changed with fresh RPMI, supplemented with 10% (v/v) fetal bovine serum (FBS) and antibiotics (100 U/ml penicillin, 100 μg/ml streptomycin) 6h post transfection. After 48h, virus-containing supernatants were harvested, filtered through 0.45-μm polyethersulfone filters (Millipore) and supplemented with 5 μg/ml polybrene (Sigma) for infection. Cells were infected by spin infection (2,000 rpm, 45 min, room temperature). 24/48h later, cells were washed and then selected with appropriate antibiotics or, in case of pINDUCER21-based vectors, sorted by FACS based on GFP signal. Selected cell populations were used for experimental investigations without further subcloning to avoid clonal effects.

### Cell lysis and western blotting

Cells were lysed in RIPA (25 mM Tris, 150 mM NaCl, 0.5% NP-40, 0.5% deoxycholate (w/v), 0.1% SDS (w/v), pH 7.4) or E1A (50 mM HEPES, 250 mM NaCl, 5 mM EDTA, 1% NP-40, pH 7.4) lysis buffer. For Native-PAGE immunoblotting, cells were lysed in a NP-40 buffer (50 mM HEPES, 150 mM NaCl, 5 mM EDTA, 10% glycerol, 1% NP-40, pH 7.4). Lysis buffers were supplemented with complete EDTA-free protease inhibitor cocktail (Roche) and Halt phosphatase inhibitor cocktail (Thermo Fisher Scientific). Lysates were cleared by centrifugation (13,000 rpm, 10 min, 4 °C), and, after protein quantification with BCA (Thermo Fisher Scientific) using BSA as standard, resolved by regular or Phos-tag-containing (20-50 μM, WAKO Chemicals) SDS–PAGE and blotted to nitrocellulose membranes (Amersham). Prior to transfer, Phos-tag SDS-PAGE gels were incubated (2 times 10 min) with transfer buffer supplemented 10 mM EDTA and then washed 10 min in transfer buffer without EDTA. For IRF5 Native-PAGE, 10ug of lysate was separated by 7.5% polyacrylamide gel without SDS. Before transfer, gels were soaked in running buffer with 0.01% SDS for 30 minutes at room temperature. After transfer, membranes were blocked by 5% non-fat dry milk in TBST and probed with indicated antibodies. In experiments in which multiple antibodies were used, equal amounts of samples were loaded on multiple SDS–PAGE gels and western blots sequentially probed with a maximum of two antibodies.

### Co-immunoprecipitation

For immunoprecipitation assays with tagged proteins, cells were lysed in E1A buffer. The appropriate amount of whole-cell lysate was used as input, and the remaining was subjected to immunoprecipitation using equilibrated CaptureSelect C-tagXL Affinity matrix beads (Thermo Fisher Scientific) overnight at 4 °C on a rotating wheel. Beads were washed three times with E1A buffer and eluted with SDS loading buffer. After quantification of the whole-cell lysate and immunoprecipitated proteins were analysed by SDS-PAGE and immunoblotting.

### PNGase F treatment

Cleared cell lysates were incubated with PNGase F (500–1,000 U per 20 μl of lysats, NEB) for 30–45 min at 37 °C. Samples were analyzed by western blotting.

### Enzyme-linked immunosorbent assay (ELISA)

All ELISA experiments were performed using diluted cell supernatants according to manufacturer’s instructions as described ^37^.

### Quantitative real-time PCR (qPCR)

Cells were collected and total RNAs were isolated using a Quick-RNA Miniprep Kit (Zymo Research). Reverse transcription was performed using RevertAid First Strand cDNA Synthesis Kit (Thermo Fisher Scientific) using oligo (dT) primers. Real-time PCR was performed using LightCycler^®^ 480 SYBR Green I Master (Roche). Gene-specific primers used are described in Table S2. Samples were analyzed on LightCycler 480 (Roche). Data were analysed and Ct values were calculated using LightCycler Software version 1.5 (Roche). Results were obtained using the 2^−ΔΔCt^ method, using HPRT1 as reference.

### Confocal microscopy

These experiments were carried out as previously described ^37^. In brief, 3.5 × 10^5^ cells were seeded in 24-well plates on coverslips and treated with 10 nM PMA (Sigma) overnight to induce adherence. Cells were washed with PBS, fixed (PBS, 2% formaldehyde (Sigma)) for 20 min, permeabilized and blocked in blocking solution (PBS, 0.3% saponin (Sigma), 10% FBS) for 1 h. Afterwards, cells were stained 1h at room temperature with the indicated primary antibodies, rabbit anti-HA (no. 3724, Cell Signaling, 1:1,000) and mouse anti-LAMP2 (sc-18822, Santa Cruz, 1:1,000) in blocking solution. After three washes of blocking solution, cells were stained with goat anti-mouse (AlexaFluor594, A11005) and anti-rabbit (AlexaFlour488, A11034) antibodies (Invitrogen, 1:1,000) for 1h at room temperature. Cells were then washed once in blocking solution. Nuclear counterstaining was performed with DAPI (Invitrogen). After three washes with blocking buffer and one wash with PBS, cover glasses were mounted onto microscope slides using ProLong Gold (Invitrogen) antifade reagent. Images were acquired on a confocal laser scanning microscope (Zeiss LSM 880, Carl Zeiss) and analysed using ZEN 2.3 (Carl Zeiss).

### Quantification and statistical analysis

### Statistical analysis

Statistical analyses and graphs were made using GraphPad Prism 9 software (GraphPad). The number of experiments or biological replicates (n) used for the statistical evaluation of each experiment is indicated in the corresponding figure legends. The data are plotted as a mean ± SD as indicated.

---

**Figure S1.**
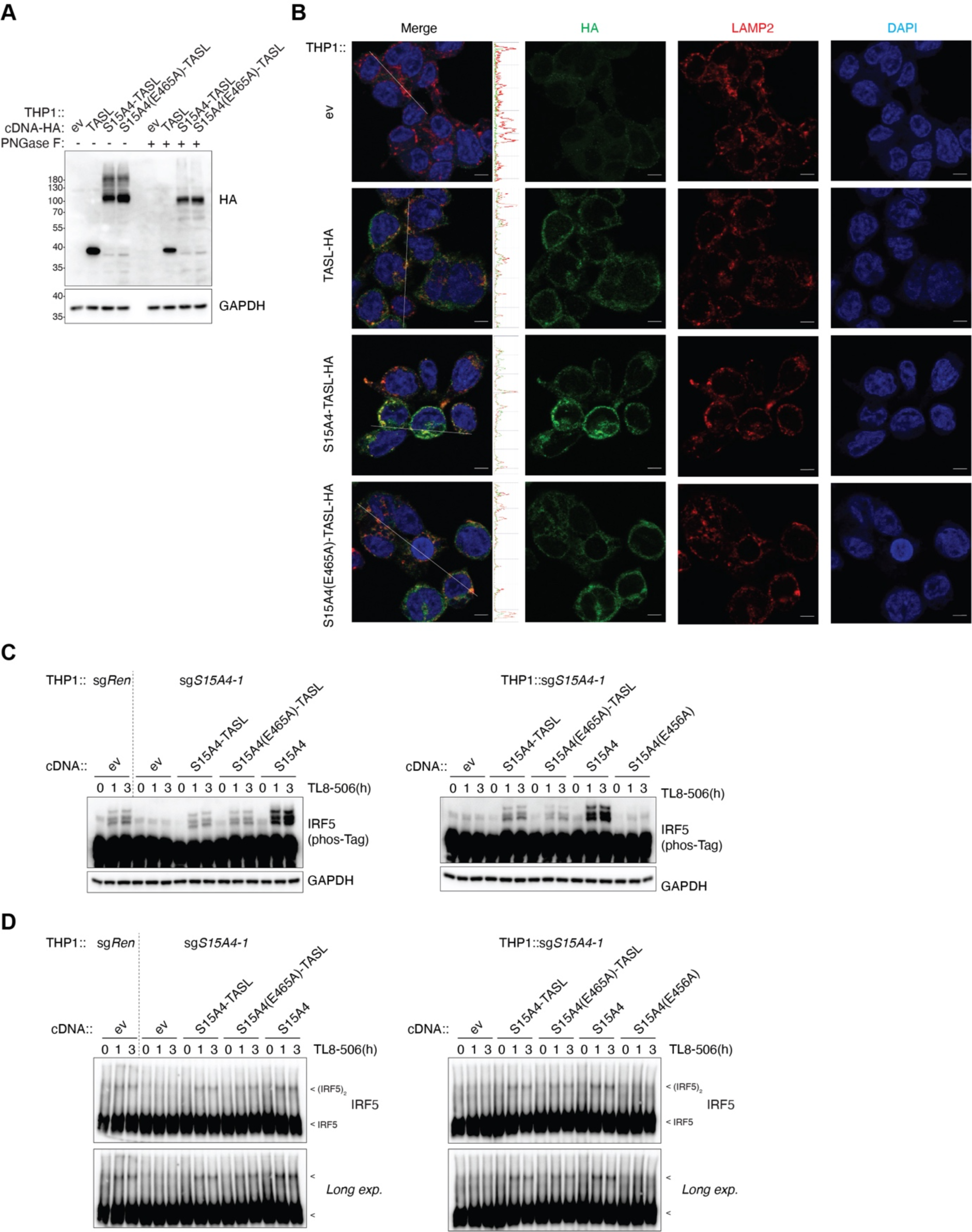
Transport-deficient SLC15A4(E465A)-TASL fusion protein localize to the endolysosomal compartment and rescues TLR8-induced IRF5 activation, related to Figure 1. (A) Immunoblots of THP1 cells stably reconstituted with indicated C-terminal HA-tagged TASL constructs. (B) Confocal microscopy of indicated THP1 cells. Green, anti-HA; red, anti-LAMP2; blue, DAPI. (C and D) Indicated THP1 cells were stimulated with TLR8 ligand TL8-506 (0.5 μg ml−1, for 0-3 h). Cell lysates were analysed by immunoblotting using Phos-tag SDS-PAGE (C) or Native PAGE (D). long exp.: long exposure. Arrows indicate IRF5 monomer or dimer. In C-D, data are representative of two independent experiments.

**Figure S2.**
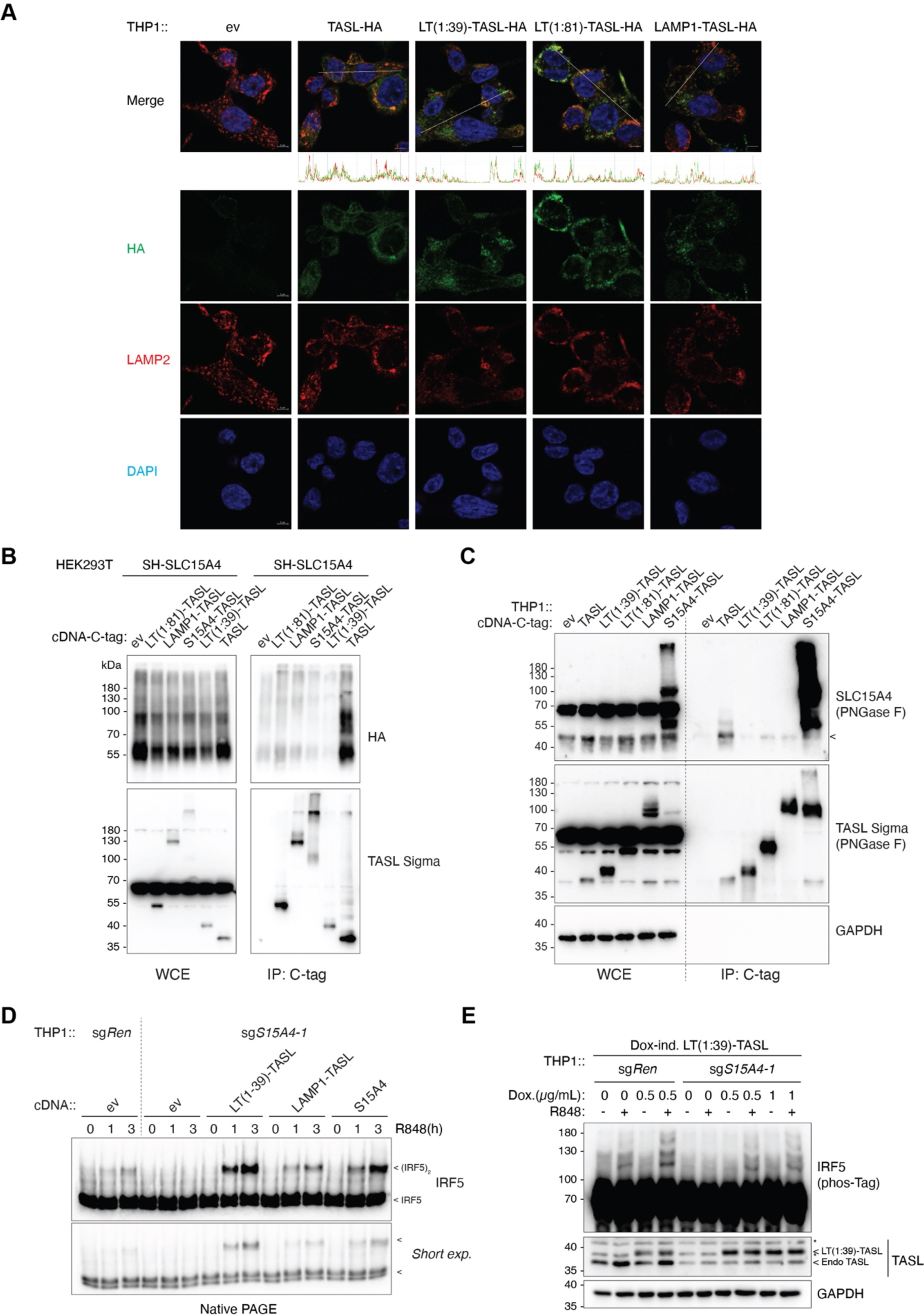
TASL fusions localize to the endolysosome and restore IRF5 activation independently of SLC15A4, related to Figure 2. (A) Confocal microscopy of indicated THP1 cells. Green: anti-HA; red: anti-LAMP2; blue: DAPI. (B and C) Immunoprecipitates (IP, C-tag) and whole cell lysates (WCE) from transiently transfected HEK293T (B) or stably transduced THP1 cells (C) were analysed by immunoblotting with indicated antibodies. Where indicated, lysates were treated with PNGase F. SH, Strep-HA tag. (D) Native PAGE immunoblots of indicated THP1 cells stimulated with R848 (5 μg ml−1, for 0-3h). short exp.: short exposure. Arrows represent IRF5 monomer or dimer. (E) Immunoblots of knockout THP1 cells reconstituted with doxycycline-inducible LAMTOR1(1:39)-TASL. 24h after the addition of doxycycline as indicated, cells were stimulated with R848 (5 μg ml−1 for 3 h). Arrows indicate LT(1:39)-TASL or endogenous TASL. In B-E, data are representative of two independent experiments.

**Figure S3.**
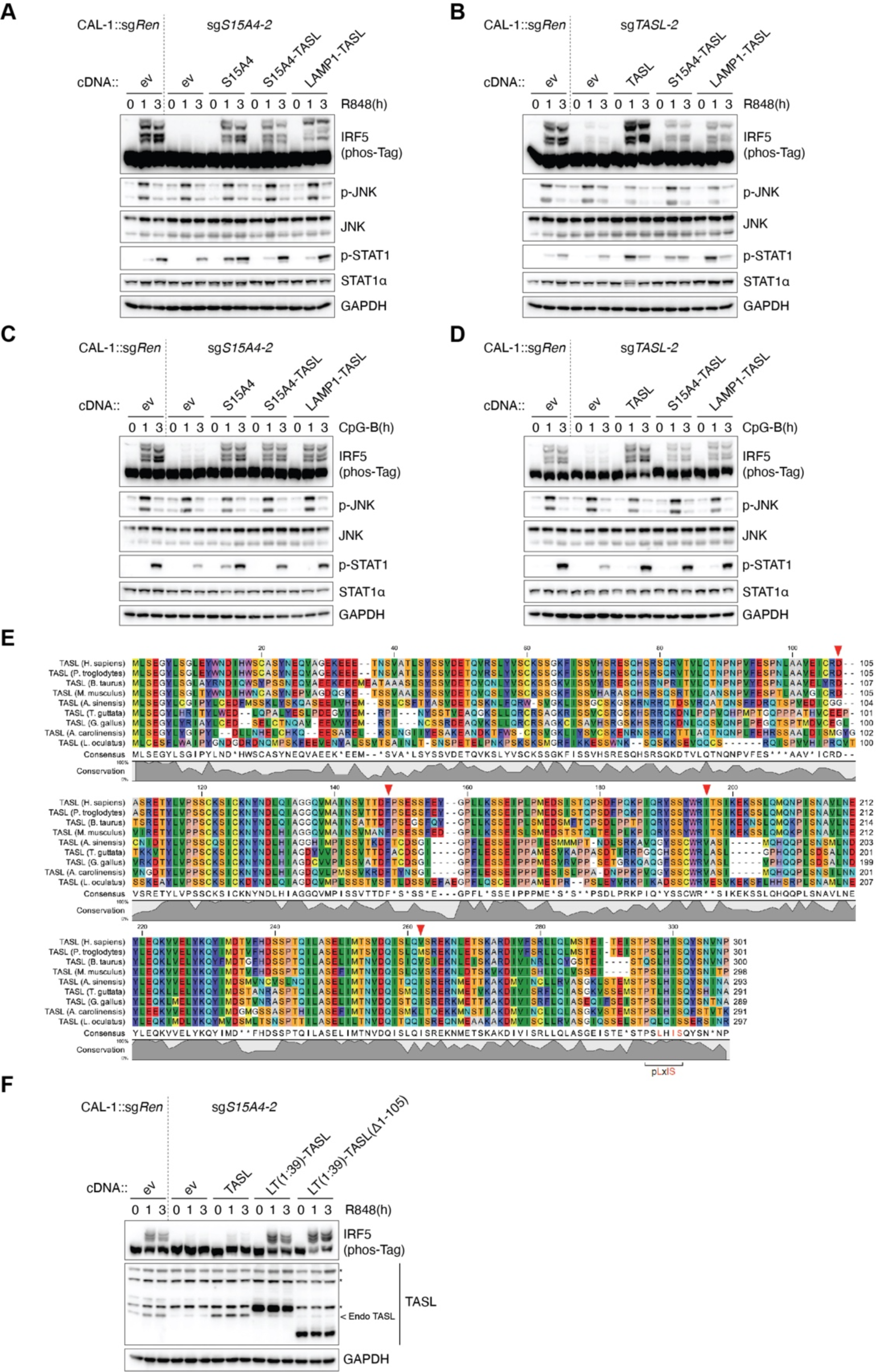
Endolysosomal TASL fusions sustains TLR7- and TLR9-induced IRF5 activation in CAL-1 pDCs independently of SLC15A4, related to Figure 3. (A-D) Immunoblot of lysates from indicated CAL-1 cells stimulated with R848 (5 μg ml−1 for 0-3 h) (A and B) or CpG-B (5 μM for 0-3 h) (C and D). (E) Multiple sequence alignment of TASL protein from representative vertebrate species. UniProt entry names: CX021_HUMAN, H2QYF9_PANTR, CX021_BOVIN, CX021_MOUSE, A0A1U7RX84_ALLSI, H0Z9M3_TAEGU, A0A1L1RS25_CHICK, G1KG99_ANOCA, W5NMP6_LEPOC. Residues colours indicate amino acid property. pLxIS motif is highlighted. Red triangles indicate the starting residues of LT(1:39)-TASL deletion constructs (related to Figure 3H-I). (F) Immunoblot of lysates from indicated CAL-1 cells stimulated with R848 (5 μg ml−1 for 0-3 h), Asterisks denote unspecific bands. In A-D, F, data are representative of two independent experiments.

**Figure S4.**
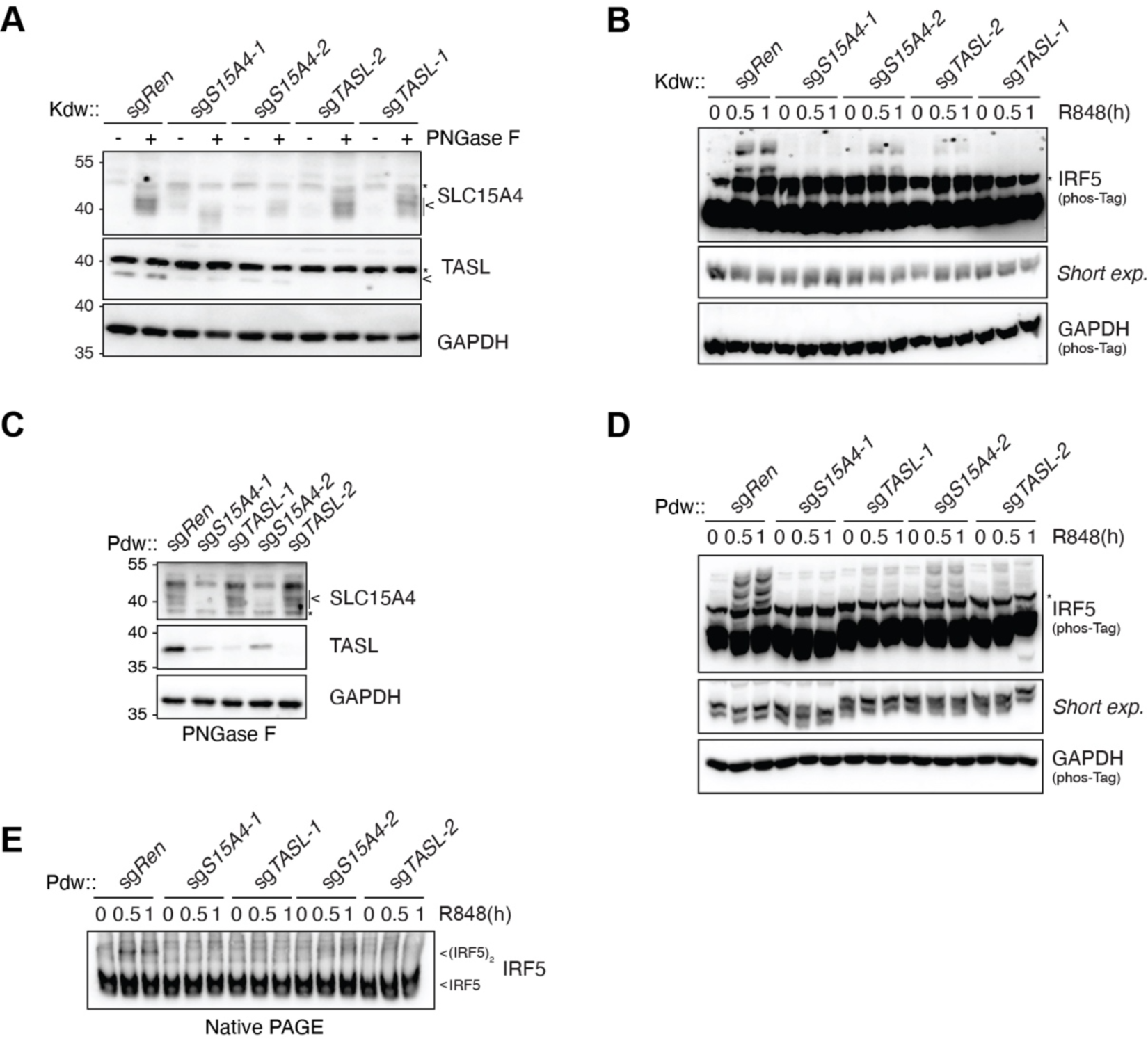
The SLC15A4-TASL complex is required for TLR7/8-induced IRF5 activation in human B cells, related to Figure 4. (A) Immunoblots of SLC15A4-or TASL-knockout Kdw cells generated using two different sgRNA targeting each gene. Lysates were treated with PNGase F as indicated. Arrow indicates endogenous TASL. (B) Immunoblots of indicated Kdw cells stimulated with R848 (5 μg ml−1, for 0-1h). (C) Immunoblots of SLC15A4-or TASL-knockout Pdw cells generated using two different sgRNA targeting each gene. Lysates were treated with PNGase F. (D) Immunoblots of knockout Pdw cells stimulated with R848 (5 μg ml−1, for 0-1h). (E) Native PAGE immunoblots in knockout Pdw cells stimulated with R848 (5 μg ml−1, for 0-1h). Arrows indicate IRF5 monomer or dimer. In A-E, data are representative of two independent experiments. Asterisks denote unspecific bands.

**Table S1.**
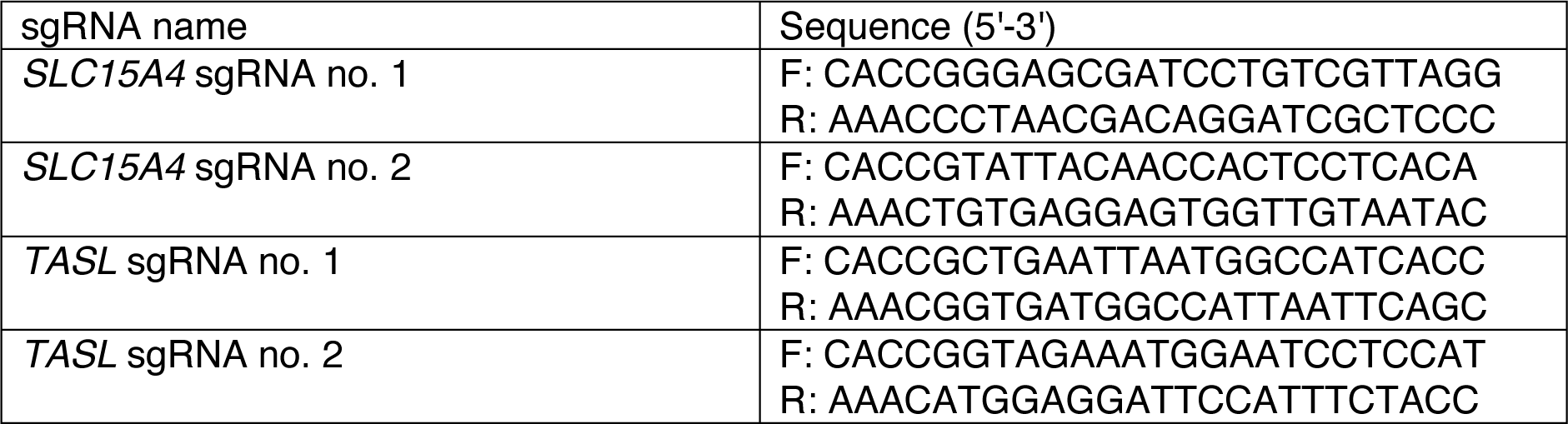
Oligonucleotides used for sgRNAs cloning in this study. Related to STAR Methods

**Table S2.**
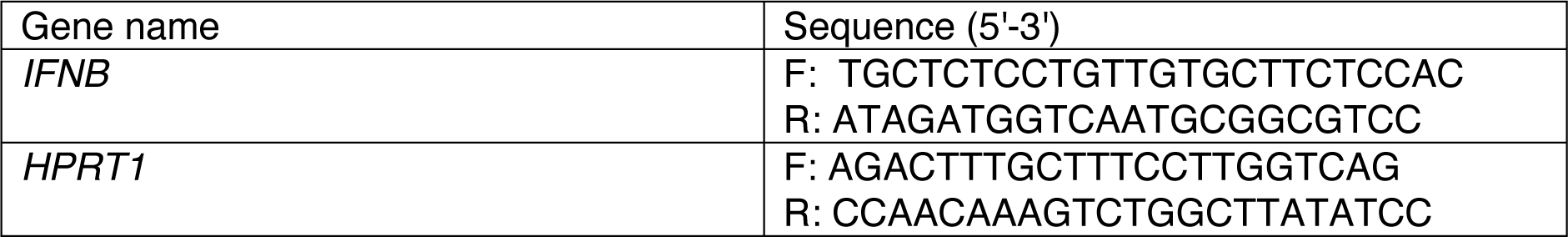
Primers used for qPCR in this study. Related to STAR Methods

